# Assessing molecular phylogenetics of Hydroptilidae (Trichoptera) subfamily lineages, with notes on biogeography

**DOI:** 10.1101/2023.09.20.558543

**Authors:** Kelly M. Murray-Stoker, Shannon J. McCauley

## Abstract

The microcaddisfly family Hydroptilidae is the most species-rich within the insect order Trichoptera and has a global distribution. The taxonomy and systematics of this group remains understudied in proportion to its diversity. Here, we present a phylogenetic assessment of subfamily relationships that includes five of the six Hydroptilidae subfamilies (Hydroptilinae, Leucotrichiinae, Ochrotrichiinae, Orthotrichiinae, and Stactobiinae) using publicly available molecular data. Our analyses recovered Leucotrichiinae, Ochrotrichiinae, and Stactobiinae as monophyletic and Hydroptilinae as paraphyletic. Based on the diversity and distribution of taxa, much more representation of species is needed to fully understand the relationships of subfamilies; however, our results suggest a need to revisit the placement of *Ithytrichia*, which concurs with other recent molecular work on Hydroptilidae. Further clarity into the relationships of Hydroptilidae will be important for understanding the variability and phylogenetic signal of ecological traits.

## Introduction

Phylogenetic analyses are essential to visualizing the processes of evolution and the generation of biodiversity. Many different types of analyses can be conducted using phylogenetic trees, such as linking of trait evolution with phylogenetic diversity. For example, in the caddisflies, studying the feeding ecology of larvae within a phylogenetic context allowed Pauls et al. (2008) to gain new insights into the relationship of the evolution of grazing within the family Limnephilidae and the high degree of species diversity. Researchers can also identify synapomorphies, understand how many times certain traits have evolved within a taxonomic group, and infer the ancestral state of a group, such as the conclusion by Thomas et al. (2020) that phylogenetic analyses of Trichoptera show evidence of ancestral larval case-building behavior. Another application of phylogenetic analyses is the incorporation of biogeographic data to determine what environmental factors may have contributed to diversification of certain groups.

Failing to study and consider phylogenetic relationships can result in taxonomy based on unnatural groupings rather than groupings with shared evolutionary history. In such cases, certain, shared traits could be incorrectly assumed to be homologous when they instead may be the result of convergent evolution. Additionally, phylogenetic analyses will allow researchers to determine whether traits are synapomorphic or plesiomorphic. Early studies of Trichoptera grouped four families, Rhyacophilidae, Glossosomatidae, Hydroptilidae, and Hydrobiosidae into a distinct suborder (“Spicipalpia”) based on shared morphological characteristics (Wiggins and Wichard, 1989). Now, however, with molecular phylogenetic techniques, Thomas et al. (2020) have determined these taxa do not form a monophyletic group, but instead are separate, basal lineages to the suborder Integripalpia (though recent analyses indicate that Hydroptilidae may actually be sister to the suborder Annulipalpia; Ge et al., 2022).

The increasing accessibility of molecular analyses over the past few decades has enabled the use of phylogenetic analyses in community ecology, allowing researchers to assess the role of evolution in community assembly (Webb et al., 2002). However, there have been critiques of such methods, particularly of the assumption that relatedness necessarily confers ecological, or trait, similarity (Cadotte et al., 2017). To understand fully how traits and relatedness correlate within important freshwater taxa like Trichoptera, in-depth study of both sides of this dynamic, phylogenetics and functional trait ecology, is needed. Here, we are interested in contributing towards the body of Trichoptera systematic knowledge so that any future investigations of functional ecology can operate under up-to-date phylogenies.

We aim to determine how molecular phylogenetic analysis can help resolve the subfamily relationships within the caddisfly family Hydroptilidae, the “microcaddisflies.” For several decades (until recent work published by Thomson et al., 2022), the most prominent family-level phylogeny of Hydroptilidae was constructed with morphological characters by Marshall (1979). Since then, many species and genera have been described within the family (Morse, 2023; Thomson, 2023). Intensive molecular phylogenetic work has been done within one subfamily, Leucotrichiinae (Santos et al., 2016), though a thorough treatment of each of the other subfamilies is still needed.

We also relate biogeography and species diversity to the phylogenetic analysis of the subfamily lineages to assess whether certain lineages are concentrated in particular parts of the world. Hydroptilidae is the most diverse family of Trichoptera, with over 2000 species and a worldwide distribution (Morse, 2023; Thomson, 2023); relating biogeographical occurrences to phylogenetic analyses may indicate spatial patterns of Hydroptilidae evolution. Here, we present preliminary analyses using publicly available data on GenBank (NCBI, 2021). The availability of molecular data does not reflect the diversity of this caddisfly family; nevertheless, the representation of most of the subfamilies will hopefully allow for some preliminary conclusions regarding within-family lineages.

## Methods

Thirty-seven caddisfly (Insecta: Trichoptera) species in the family Hydroptilidae were chosen for these analyses based on the availability of public data, representing 22 genera and five out of the six described subfamilies in Hydroptilidae (Table 1). We selected all species which had sequences for at least two out of the following three loci available on GenBank as of March 2021 (NCBI, 2021): mitochondrial cytochrome oxidase subunit I (COI, ∼650 bp), nuclear elongation factor 1 alpha (EF-1α, ∼350 bp), and nuclear carbomoylphosphate synthase (CAD, ∼850 bp). Based on these criteria, we did not have representation of the subfamily Neotrichiinae. We chose *Glossosoma nigrior* Banks 1911 (Glossosomatidae) as an outgroup taxon. We also retrieved data on the number of species and geographic records represented within each genus studied here in order to compare diversity and distribution among subfamilies; data were retrieved from the Trichoptera World Checklist (Morse, 2023). We estimated phylogenies using both Maximum Likelihood (ML) and Bayesian Inference approaches.

**Table 1.**
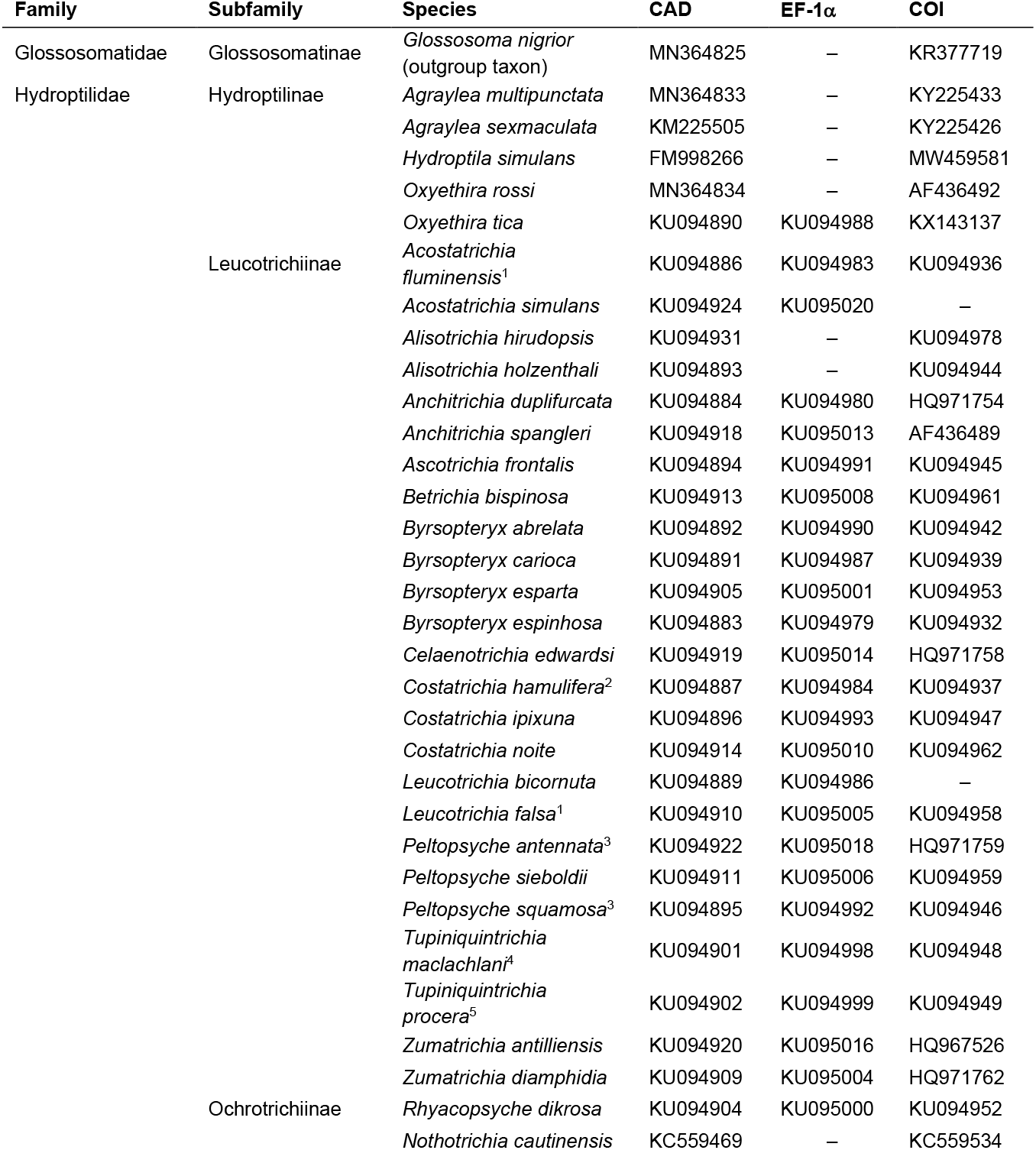
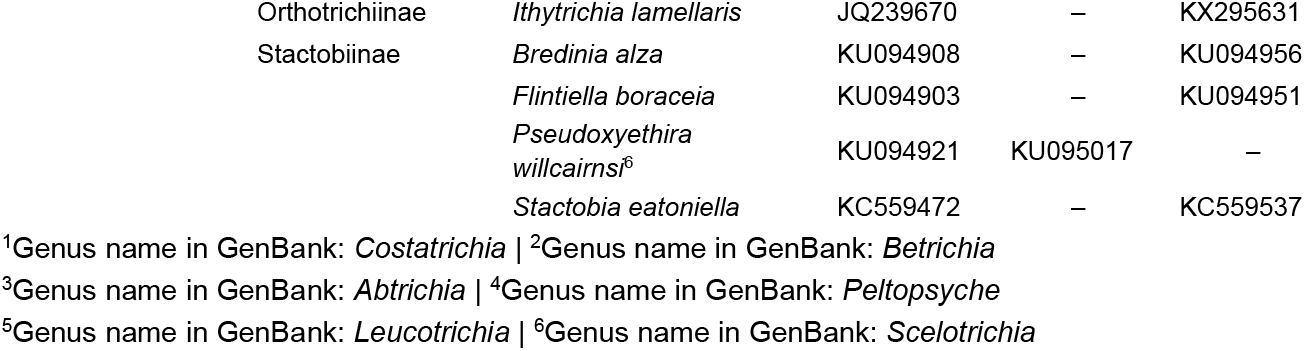
Hydroptilidae species (plus Glossosomatidae outgroup) used in phylogenetic analyses. All sequences for the three loci of interest (CAD, EF-1α, and COI) were obtained from GenBank (NCBI, 2021); accession numbers are listed. Species names that have been updated in recent taxonomic literature are listed with the current name, and the original names in GenBank are noted below the table.

| Family | Subfamily | Species | CAD | EF-1α | COI |
| --- | --- | --- | --- | --- | --- |
| Glossosomatidae | Glossosomatinae | *Glossosoma nigrior*<br>(outgroup taxon) | MN364825 | – | KR377719 |
| Hydroptilidae | Hydroptilinae | *Agraylea multipunctata* | MN364833 | – | KY225433 |
|  |  | *Agraylea sexmaculata* | KM225505 | – | KY225426 |
|  |  | *Hydroptila simulans* | FM998266 | – | MW459581 |
|  |  | *Oxyethira rossi* | MN364834 | – | AF436492 |
|  |  | *Oxyethira tica* | KU094890 | KU094988 | KX143137 |
|  | Leucotrichiinae | *Acostatrichia fluminensis*<sup>1</sup> | KU094886 | KU094983 | KU094936 |
|  |  | *Acostatrichia simulans* | KU094924 | KU095020 | – |
|  |  | *Alisotrichia hirudopsis* | KU094931 | – | KU094978 |
|  |  | *Alisotrichia holzenthali* | KU094893 | – | KU094944 |
|  |  | *Anchitrichia duplifurcata* | KU094884 | KU094980 | HQ971754 |
|  |  | *Anchitrichia spangleri* | KU094918 | KU095013 | AF436489 |
|  |  | *Ascotrichia frontalis* | KU094894 | KU094991 | KU094945 |
|  |  | *Betrichia bispinosa* | KU094913 | KU095008 | KU094961 |
|  |  | *Byrsoteryx abrelata* | KU094892 | KU094990 | KU094942 |
|  |  | *Byrsoteryx carioca* | KU094891 | KU094987 | KU094939 |
|  |  | *Byrsoteryx esparta* | KU094905 | KU095001 | KU094953 |
|  |  | *Byrsoteryx espinhosa* | KU094883 | KU094979 | KU094932 |
|  |  | *Celaenotrichia edwardsi* | KU094919 | KU095014 | HQ971758 |
|  |  | *Costatrichia hamulifera*<sup>2</sup> | KU094887 | KU094984 | KU094937 |
|  |  | *Costatrichia ipixuna* | KU094896 | KU094993 | KU094947 |
|  |  | *Costatrichia noite* | KU094914 | KU095010 | KU094962 |
|  |  | *Leucotrichia bicornuta* | KU094889 | KU094986 | – |
|  |  | *Leucotrichia falsa*<sup>1</sup> | KU094910 | KU095005 | KU094958 |
|  |  | *Peltopsyche antennata*<sup>3</sup> | KU094922 | KU095018 | HQ971759 |
|  |  | *Peltopsyche sieboldii* | KU094911 | KU095006 | KU094959 |
|  |  | *Peltopsyche squamosa*<sup>3</sup> | KU094895 | KU094992 | KU094946 |
|  |  | *Tupiniquintrichia maclachlan*<sup>4</sup> | KU094901 | KU094998 | KU094948 |
|  |  | *Tupiniquintrichia procera*<sup>5</sup> | KU094902 | KU094999 | KU094949 |
|  |  | *Zumatrichia antilliensis* | KU094920 | KU095016 | HQ967526 |
|  |  | *Zumatrichia diamphidia* | KU094909 | KU095004 | HQ971762 |
|  | Ochrotrichiinae | *Rhyacopsyche dikrosa* | KU094904 | KU095000 | KU094952 |
|  |  | *Nothotrichia cautinensis* | KC559469 | – | KC559534 |

|  |  |  |  |  |
| --- | --- | --- | --- | --- |
| Orthotrichiinae | *Ithytrichia lamellaris* | JQ239670 | – | KX295631 |
| Stactobiinae | *Bredinia alza* | KU094908 | – | KU094956 |
|  | *Flintiella boracea* | KU094903 | – | KU094951 |
|  | *Pseudoxyethira willcairnsi*<sup>6</sup> | KU094921 | KU095017 | – |
|  | *Stactobia eatoniella* | KC559472 | – | KC559537 |

<sup>1</sup>Genus name in GenBank: *Costatrichia* | <sup>2</sup>Genus name in GenBank: *Betrichia*
<sup>3</sup>Genus name in GenBank: *Abtrichia* | <sup>4</sup>Genus name in GenBank: *Peltopsyche*
<sup>5</sup>Genus name in GenBank: *Leucotrichia* | <sup>6</sup>Genus name in GenBank: *Scelotrichia*

Sequence alignment was completed in MUSCLE v3.8.31 (Edgar, 2004), and sequences were concatenated in Mesquite v3.61 (Maddison and Maddison, 2019). For each optimality criterion, we used both a dataset that was a combination of nucleotide sequences for the CAD and EF-1α loci and the amino acid sequences of the COI locus. we used the amino acid translations of COI sequences available on GenBank (NCBI, 2021) for each species. The data were partitioned by codon position for each locus.

Model testing and Maximum Likelihood analyses were conducted in IQ-TREE v1.6.12 (Chernomor et al., 2016; Kalyaanamoorthy et al., 2017; Nguyen et al., 2015). Model testing was restricted to only models usable in MrBayes software (i.e., Jukes-Cantor, Felsenstein81, Hasegawa-Kishino-Yano, Kimura Two-Parameter, General Time Reversible, Symmetric), and the partitioned codons were merged into the minimum number of partitions possible. Each tree was constructed with 1000 replications of a heuristic search and 1000 ultrafast bootstrap replicates.

Bayesian inference analyses were conducted in MrBayes v3.2.7a (Huelsenbeck and Ronquist, 2001; Ronquist et al., 2012; Ronquist and Huelsenbeck, 2003). Models for each partition were specified from the results of model testing in IQ-TREE. We used flat priors for each Bayesian analysis and allowed nucleotide substitution to vary within each partition, with two runs per simulation and a burn-in fraction of 0.25. Markov chain Monte Carlo runs generated by MrBayes were evaluated using Tracer v1.7.1 (Rambaut et al., 2018). The simulation was continued until all or most of the effective sample sizes of each parameter were greater than or equal to 200 and the average standard deviation of split frequencies was less than 0.01 (4,000,000 generations), indicating adequate convergence of the two runs. All trees were visualized using the software FigTree v1.4.4 (Rambaut et al., 2018); subfamily designations were added, and text was edited for species name formatting and readability of node labels in Adobe Illustrator v.25.2.1 (Adobe Inc, 2021).

## Results

The final partitions used for both the ML and Bayesian analyses, as well as the best models for each partition for the combined nucleotide-amino acid dataset, are listed in Table 2. Partitions and models were similar between the datasets for the CAD and EF-1α loci, both of which included a mixture of General Time Reversible (Tavaré, 1986) and Hasegawa-Kishino-Yano models (Hasegawa et al., 1985).

**Table 2.** Models used for the partitions of the combined nucleotide-amino acid data set. Best models for each partition were identified using IQ-TREE v1.6.12 (Chernomor et al., 2016; Kalyaanamoorthy et al., 2017; Nguyen et al., 2015). Meaning of the abbreviations are as follows. GTR: General Time Reversible (nucleotide model, unequal rates, and unequal base frequencies), HKY: Hasegawa-Kishino-Yana (nucleotide model, unequal transition/transversion rates and unequal base frequencies), mtREV (amino acid model for vertebrate mitochondrial loci), +F: indicating empirical base frequencies, +I: allowing for invariable sites, +G4: Gamma model with 4 rate categories.

| Partition (Locus + codon position) | Model |
| --- | --- |
| COI | mtREV + I + G4 |
| CAD1 | GTR + F + I + G4 |
| CAD2_EF1α2 | GTR + F + I + G4 |
| CAD3_EF1α3 | HKY + F + I + G4 |
| EF1α1 | GTR + F + G4 |

Both analyses resulted in roughly similar trees (Figure 1, Figure 2), for example, recovering Leucotrichiinae, Ochrotrichiinae, and Stactobiinae as monophyletic and Hydroptilinae as paraphyletic. The trees showed Stactobiinae as a sister clade to the rest of the subfamily lineages and Ochrotrichiinae as the sister group to Leucotrichiinae (Figure 1, Figure 2). However, the ML analysis (Figure 1, log likelihood: –15648.990) showed relatively low percent support for the node separating Ochrotrichiinae + Leucotrichiinae from the rest of the Hydroptilidae, with 48% bootstrap support. This node also had relatively low posterior probability as determined by the Bayesian analyses (0.6988, Figure 2).

**Figure 1.**
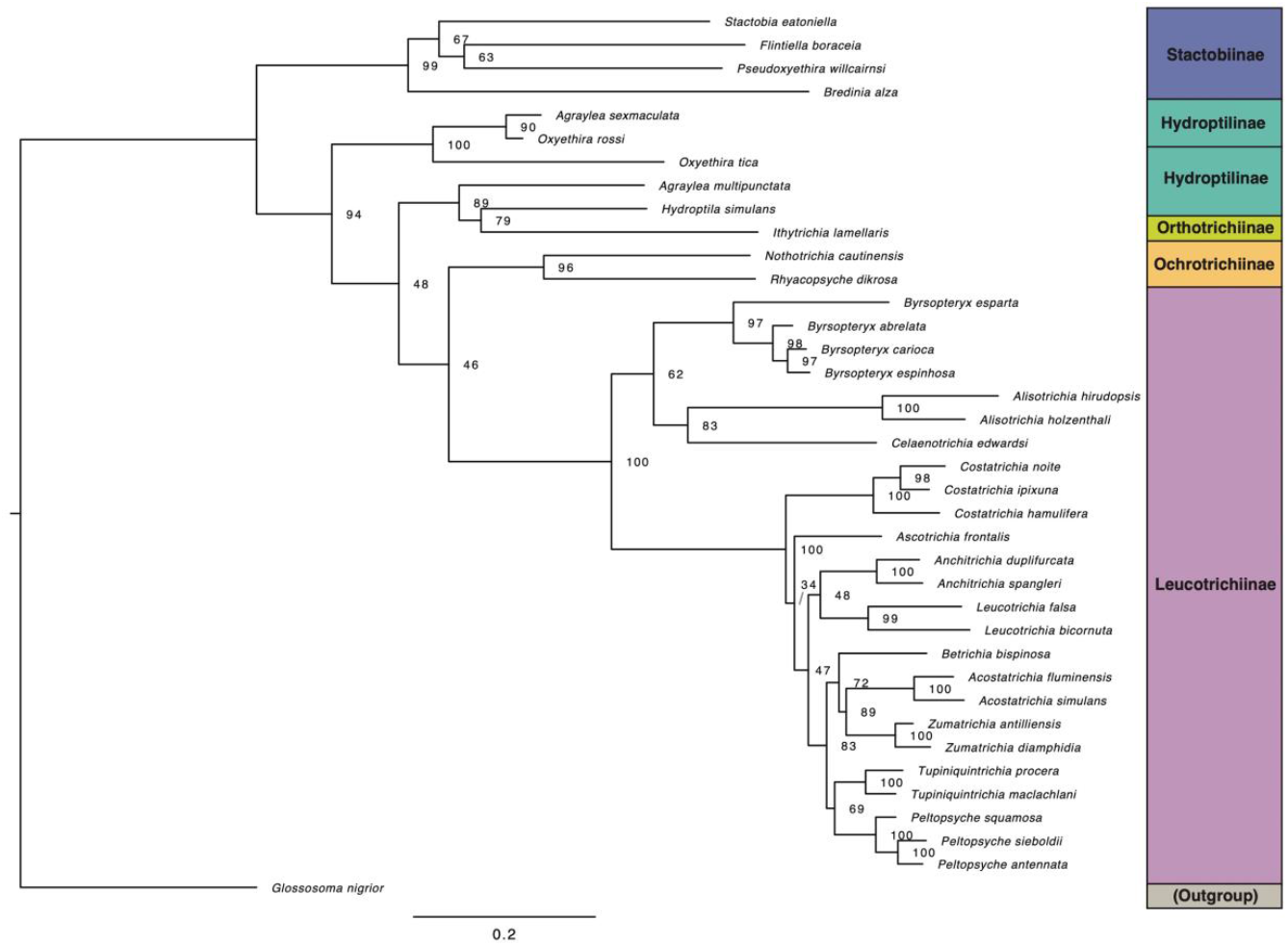
Maximum likelihood tree for the mixed dataset of CAD and EF-1α nucleotide sequences and COI amino acid sequences of Hydroptilidae species; log likelihood: –15648.990. Subfamily membership is denoted by the color-coded key; split subfamily designation between adjacent species is to denote non-monophyly of that subfamily (i.e., Hydroptilinae). Species names are updated from GenBank designations to reflect current taxonomic nomenclature. Node labels indicate bootstrap support percentage. Scale: number of substitutions per site.

**Figure 2.**
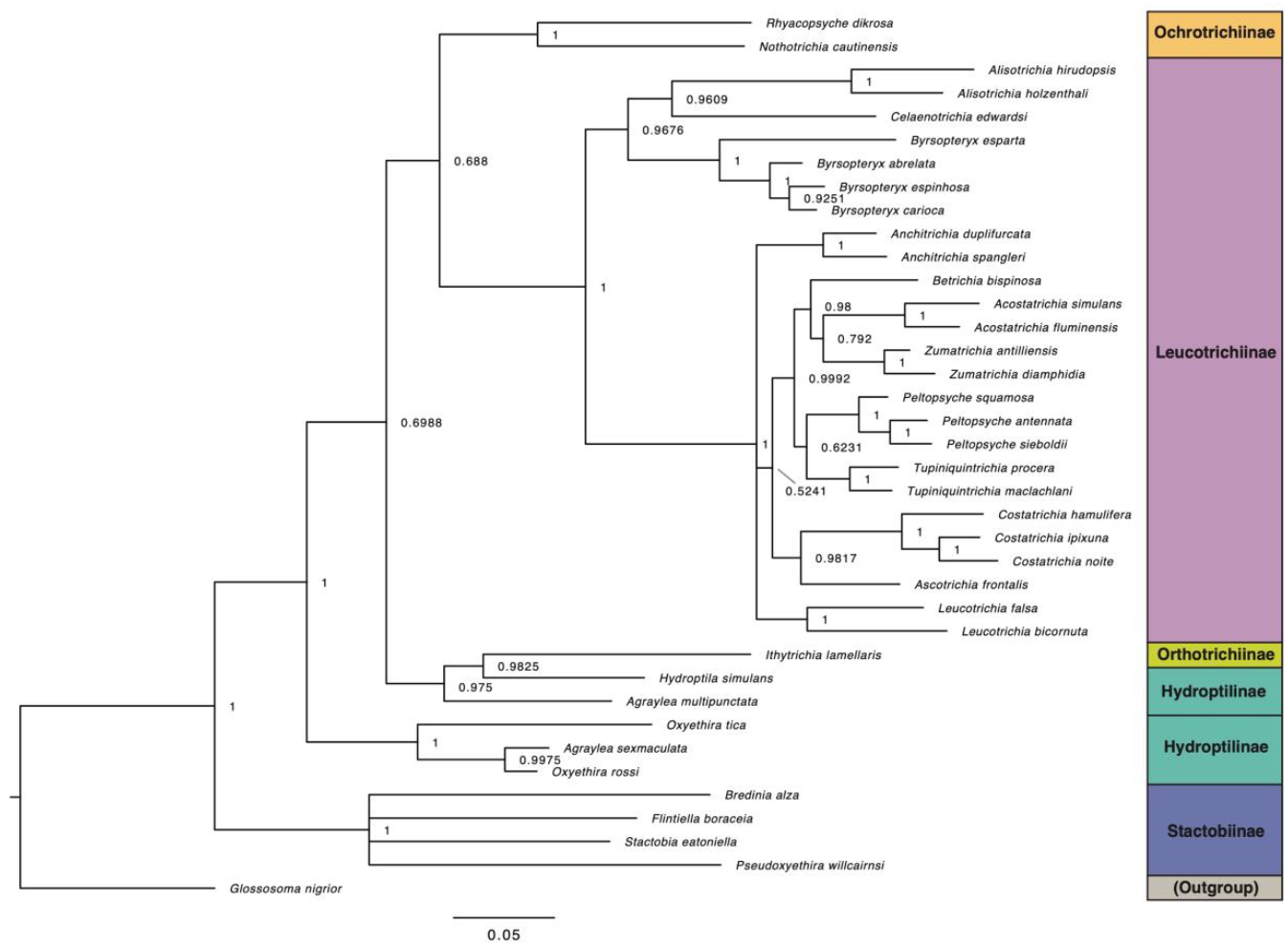
Bayesian inference tree for the mixed dataset of CAD and EF-1α nucleotide sequences and COI amino acid sequences of Hydroptilidae species. Subfamily membership is denoted by the color-coded key; split subfamily designation between adjacent species is to denote non-monophyly of that subfamily (i.e., Hydroptilinae). Species names are updated from GenBank designations to reflect current taxonomic nomenclature. Node labels indicate posterior probability. Scale: number of substitutions per site.

In terms of the paraphyly of Hydroptilinae, both trees showed *Hydroptila simulans* Mosely 1920 (Hydroptilinae) and *Ithytrichia lamellaris* Eaton 1873 (Orthotrichiinae) as sister taxa, with 79% for the ML tree, (Figure 1), though relatively high posterior probability for the Bayesian tree (0.9825, Figure 2). In both trees, *Agraylea multipunctata* Curtis 1834 (Hydroptilinae) was recovered as the sister taxon to the *Hydroptila simulans* + *Ithytrichia lamellaris* clade, though its congener *Agraylea sexmaculata* Curtis 1834 was placed in a clade with two species of *Oxyethira* (Hydroptilinae).

There was variability in the diversity and distributions of the genera examined in this study (Table 3). The leucotrichiin species are mostly confined to the Neotropical biogeographic region, with some representation in the Nearctic. Genera within Leucotrichiinae ranged in total species number from 1 to 61 (Table 3). By contrast, two of the three genera within the subfamily Hydroptilinae included in the analyses (*Hydroptila* and *Oxyethira*) both have recorded distributions in seven biogeographic regions and contain more than 250 species each (Table 3). Stactobiinae also has relatively widespread distribution, and the genus *Stactobia* contains 164 species (Table 3).

**Table 3.** Global distribution and diversity of the Hydroptilidae genera studied here, listed by biogeographic region: Afrotropic (AF), Australasia-Oceania (AU), East Palearctic (EP), Indomalaya (IM), Nearctic (NA), Neotropic (NT), and West Palearctic (WP). These occurrences apply to the whole genus, rather than just the taxa studied here; also listed are the total number of species in each genus. Distributional and diversity data retrieved from the Trichoptera World Checklist (Morse, 2023) and from Thomson (2023).

| Subfamily | Genus | Regional occurrences | Number of species |
| --- | --- | --- | --- |
| Hydroptilinae | *Agraylea* | EP, NA, IM, WP | 8 |
|  | *Hydroptila* | AT, AU, EP, IM, NA, NT, WP | 495 |
|  | *Oxyethira* | AT, AU, EP, IM, NA, NT, WP | 256 |
| Leucotrichiinae | *Acostatrichia* | NT | 15 |
|  | *Alisotrichia* | NA, NT | 61 |
|  | *Anchitrichia* | NA, NT | 8 |
|  | *Ascotrichia* | NT | 6 |
|  | *Betrichia* | NT | 10 |
|  | *Byrsoteryx* | NT | 16 |
|  | *Celaenotrichia* | NT | 1 |
|  | *Costatrichia* | NT | 20 |
|  | *Leucotrichia* | NT | 45 |
|  | *Peltopsyche* | NT | 6 |
|  | *Tupiniquintrichia* | NT | 2 |
|  | *Zumatrichia* | NA, NT | 53 |
| Ochrotrichiinae | *Nothotrichia* | NA, NT | 6 |
|  | *Rhyacopsyche* | NT | 30 |
| Orthotrichiinae | *Ithytrichia* | EP, NA, NT, WP | 7 |
| Stactobiinae | *Bredinia* | NA, NT | 17 |
|  | *Flintiella* | NT | 17 |
|  | *Pseudoxyethira* | AT, AU, EP, IM | 64 |
|  | *Stactobia* | AT, EP, IM, WP | 164 |

## Discussion

The general similarity of the tree topologies from the ML and Bayesian Inference analyses lends relative confidence in these results. Analyzing the species richness and biogeography of the taxa included in the phylogenetic analyses showed that the amount of molecular study has been unequally distributed throughout Hydroptilidae both taxonomically and geographically. Revision of the Hydroptilinae and Orthotrichiinae subfamilies and certain genera therein will likely be needed once more molecular data have been collected to reflect the high diversity of the groups. While perhaps we can infer from these results that Leucotrichiinae has diversified in the Neotropics, both the lack of available molecular data and the widespread nature of genera in other subfamilies prevents any biogeographical conclusions.

As a note, here, we are using the subfamily grouping scheme suggested by Flint (1970) and revived by Malicky (2001), while other researchers refer to these groupings as tribes, as suggested by Marshall (1979). The difference hinges on whether the group Ptilocolepidae (or Ptilocolepinae) represents a subfamily within Hydroptilidae or a separate family within Trichoptera, which has not yet been satisfactorily resolved due to a lack of data (Thomas et al., 2020). Thus, again, the continuation of molecular sampling may be helpful in progressing the understanding of Hydroptilidae and Trichoptera systematics.

The first molecular phylogeny of Hydroptilidae using both Sanger sequencing of the COI and 28S genes and targeted enrichment of ∼300 genes was recently published by Thomson et al. (2022), marking an important development since Marshall’s (1979) work, and contains more species than assessed here. Similarities with our results include a monophyletic Leucotrichiinae and an *Ithytrichia* sp. (Orthotrichiinae) recovered within Hydroptilinae. In contrast to our results, Thomson et al. (2022) recovered Ochrotrichiinae was sister to the rest of the hydroptilid subfamilies, while our results suggested Stactobiinae as sister to the other subfamily lineages, which aligned with the conclusions by Marshall (1979) using morphological data.

Taxonomic work in Hydroptilidae is typically based on adult male morphology. Recent work collecting molecular data and revising the Leucotrichiinae genera by Santos et al. (2016) and Santos (2020) has resulted in greater clarity in this part of the Hydroptilidae tree. Additionally, work by Parys and Harris (2013) showed that larval morphology can be useful in determining phylogenetic placement of genera by determining that *Nothotrichia* should be moved from Orthotrichiinae to Ochrotrichiinae, which my analyses agree with, given the molecular data available. However, Thomson et al. (2022) found that *Nothotrichia* did not group with the rest of the Ochrotrichiinae. Implementing diverse approaches to taxonomic and systematic research is important; my analyses here did not incorporate morphological traits, though both adult and larval morphology could be incorporated into future studies of lineages within Hydroptilidae.

Systematics research is of course important in its own right but achieving further clarity of the phylogenetic relationships of taxa can also be important to other fields of study. In ecological studies, for example the biomonitoring of streams to assess freshwater health based on invertebrate fauna, not enough research has been done (or is even possible) to know species-specific traits and pollution tolerance values in diverse groups like Trichoptera (Poff et al., 2006). Therefore, taxa within the same genus are often assumed to have the same tolerance values. This may or may not be the case (Cadotte et al., 2017; Resh and Unzicker, 1975), though phylogenetic analyses can at least determine whether taxonomy adequately reflects evolutionary relationships. For example, an assumption of degree of relatedness may be more accurate regarding two *Costatrichia* species (Leucotrichiinae) than regarding two *Agraylea* species (Hydroptilinae). Focusing on one Trichoptera family, we have attempted to identify what inferences we can currently make about within-group relationships and what areas of the tree may need more attention through the collection of more data.

## Acknowledgements

KMS was supported by the University of Toronto Department of Ecology and Evolutionary Biology, a Connaught Graduate Fellowship, and an Ontario Graduate Scholarship. The research was also supported by the University of Toronto Mississauga Department of Biology, and the Natural Sciences and Engineering Research Council of Canada (NSERC), [RGPIN-2019-06484] to SJM. Thank you to D. de Carle for all the guidance in software use and data analysis. We are grateful to S. Kvist for initial guidance on data selection and helpful comments on an earlier draft.

